# The RNA export factor Mex67 functions as a mobile nucleoporin

**DOI:** 10.1101/789818

**Authors:** Carina Patrizia Derrer, Roberta Mancini, Pascal Vallotton, Sébastien Huet, Karsten Weis, Elisa Dultz

## Abstract

The RNA export factor Mex67 is essential for the transport of mRNA through the nuclear pore complex (NPC) in yeast, but the molecular mechanism of this export process remains poorly understood. Here, we use quantitative fluorescence microscopy techniques in live budding yeast cells to investigate how Mex67 facilitates mRNA export. We show that Mex67 exhibits little interaction with mRNA in the nucleus and localizes to the NPC independently of mRNA, occupying a set of binding sites offered by FG repeats in the NPC. The ATPase Dbp5, which is thought to remove Mex67 from transcripts, does not affect the interaction of Mex67 with the NPC. Strikingly, we find that the essential function of Mex67 is spatially restricted to the NPC since a fusion of Mex67 to the nucleoporin Nup116 rescues a deletion of MEX67. Thus, Mex67 functions as a mobile nuclear pore component, which receives mRNA export substrates in the central channel of the NPC to facilitate their translocation to the cytoplasm.

## Introduction

Directional export of mRNAs from the nucleus to the cytoplasm is a crucial step for gene expression in every eukaryotic cell. Transport is mediated by transport factors that guide the mRNA through nuclear pore complexes (NPCs), which are highly specialized transport channels embedded in the nuclear envelope. The mRNA export machinery has been characterized in multiple species and many of its constituents are highly conserved (reviewed in Heinrich et al., 2017; Kohler and Hurt, 2007).

Export of most mRNAs depends on the transport receptors Mex67-Mtr2 (Santos-Rosa et al., 1998; Strasser et al., 2000; Weis, 2002), which are also involved in the export of ribosomal subunits (Faza et al., 2012; Yao et al., 2007). Mex67 (TAP/NXF1 in Metazoa) in complex with Mtr2 mediates transport across the NPC via interactions with several nucleoporins that contain repetitive domains enriched in phenylalanine (F) and glycine (G) (Strasser et al., 2000; Terry and Wente, 2007). These FG repeats form a meshwork in the central channel of the NPC and enable selective access for shuttling nuclear transport receptors carrying different cargo between the nucleus and the cytoplasm. At steady state, Mex67 is mainly localized at the NPC (Katahira et al., 1999; Kohler and Hurt, 2007; Rodriguez et al., 2004; Strasser et al., 2000; Terry and Wente, 2007). However, it was shown to shuttle between the nucleus and cytoplasm (Segref et al., 1997). Reminiscent of the mechanism of protein transport across the NPC, the current model for mRNA export therefore postulates that Mex67-Mtr2 binds to a mature mRNP substrate in the nucleoplasm and then chaperones the mRNP through the NPC channel via successive interactions with distinct FG-repeat nuclear pore proteins (Nups) (Hautbergue et al., 2008; Stewart, 2007; Strasser et al., 2000; Strasser and Hurt, 2001).

Once the mRNA reaches the cytoplasmic side of the NPC, the final steps in mRNA export depend on two Nups localized to the cytoplasmic face of the NPC, Nup159 and Nup42, which recruit two essential mRNA export factors, the DEAD-box ATPase Dbp5 and its ATPase activator Gle1 (Adams et al., 2017; Hodge et al., 2011; Noble et al., 2011; Weirich et al., 2006). Dbp5 is proposed to impart directionality to the export process by remodeling the mRNPs emerging from the NPC in an ATP-dependent manner and displacing export factors like Mex67-Mtr2 (Cole and Scarcelli, 2006b; Folkmann et al., 2011; Liker et al., 2000; Lund and Guthrie, 2005; Montpetit et al., 2011; Snay-Hodge et al., 1998; Tran et al., 2007; Weirich et al., 2006). Disassembly of the mRNP on the cytoplasmic side of the NPC would prevent its return to the nucleus, resulting in unidirectional mRNA translocation. In addition, the release of mRNA export factors including Mex67-Mtr2 would allow them to return to the nucleus where they could function in additional rounds of mRNA export (Stewart, 2007). However, direct evidence of such a shuttling mRNA export model is lacking and it is unclear how Mex67 and Dbp5 function in mRNA export.

In order to better understand the function of Mex67 and its interplay with the nuclear pore complex and the export factor Dbp5, we used advanced fluorescence microscopy techniques including fluorescence recovery after photobleaching (FRAP), fluorescence correlations spectroscopy (FCS) and fluorescent intensity measurements. This allowed us to quantitatively characterize the interactions of Mex67 at the NPC. In line with recent observations in mammalian cells (Ben-Yishay et al., 2019), we show that the interaction of Mex67 at the NPC does not depend on cargo and is not modulated by the activity of Dbp5. Furthermore, we uncovered that Mex67 does not have to leave the NPC to mediate mRNA export. This suggests that Mex67 behave similar to a mobile nucleoporin.

## Results

### The majority of Mex67 is not bound to mRNA in the nucleus

The binding of mRNA to its transport receptor Mex67, a crucial step in mRNA export, has been proposed to occur in the nucleoplasm (Dieppois et al., 2006; Gilbert and Guthrie, 2004; Strasser and Hurt, 2001). We reasoned that mRNA binding could be detected by assessing the diffusion of Mex67, as free proteins diffuse faster than large mRNP particles. GFP, for example, exhibits an apparent diffusion coefficient of 11 μm^2^/s in the yeast nucleus (Slaughter et al., 2007), while diffusion coefficients reported for mRNAs are in the range of 0.15-0.74 μm^2^/s (Wu et al., 2012). To measure the diffusion of Mex67 and Dbp5 in living cells, we used fluorescence correlation spectroscopy (FCS), which allows to determine the mean duration required for a fluorescent molecule to diffuse through the confocal volume (termed diffusion time).

First, we measured the diffusion of control proteins. EGFP exhibited the expected fast diffusion behavior with diffusion times of 0.6 ms in both the nucleus and the cytoplasm (Figure 1B). In contrast, Gbp2, a poly(A+) RNA-binding protein that acts as surveillance factors for the selective export of spliced mRNAs exhibited a median diffusion time of 17.9 ms in the nucleus (Figure 1C), corresponding to an apparent diffusion coefficient of 0.6 μm^2^/s, which is consistent with mRNA diffusion and indicates that the majority of Gbp2 molecules are bound to mRNPs or other slowly moving complexes. Next we assessed the diffusion of Mex67 and Dbp5 in the nucleus and the cytoplasm. Dbp5 exhibited diffusion times of 1.4 ms and 1.3 ms, respectively (Figure 1B), similar to EGFP. Thus, in both compartments the majority of Dbp5 is in a rapidly diffusing, unbound state. This is consistent with the model that Dbp5 binds mRNA predominantly at the NPC, where it functions to remodel it at the cytoplasmic filaments during late steps of mRNA export (Tran et al., 2007; von Moeller et al., 2009; Zhao et al., 2002).

**Figure 1:**
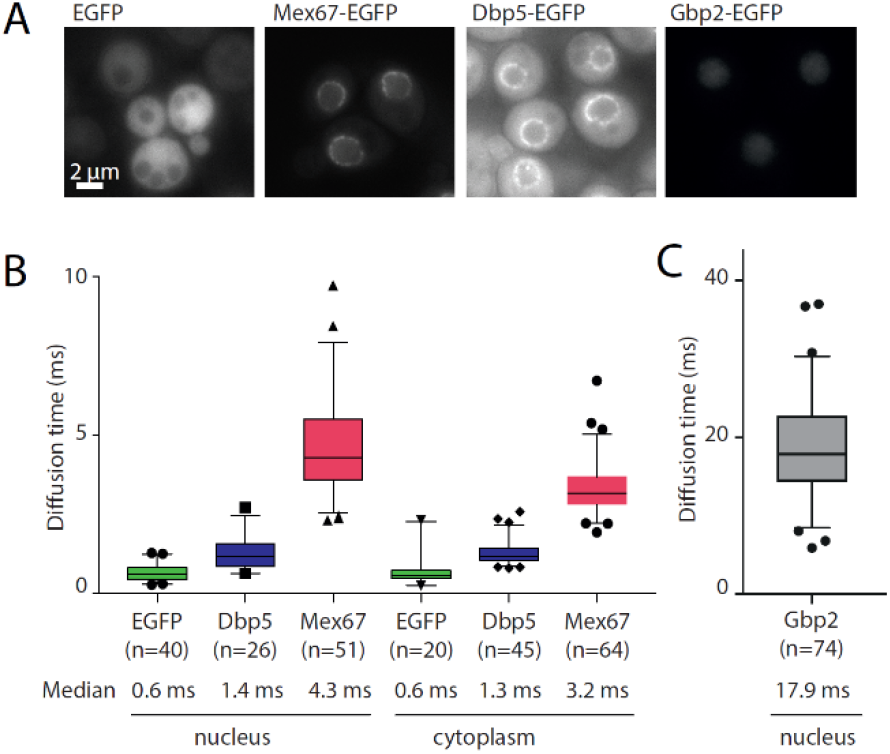
Diffusion of mRNA export factors measured by FCS. (A) Representative fluorescence wide-field microscopy images of yeast strains expressing GFP-tagged mRNA export factors. Scale bar 2 μm. (B-C) Diffusion times measured by FCS for the indicated GFP-fusion proteins. Data are pooled from three independent biological replicates. In the boxplots, the line represents the median, box represents 25-75 percentile, whiskers extend to 5-95 percentile, black symbols represent data points outside this range.

The diffusion times of Mex67-EGFP in the nucleus and cytoplasm were 4.3 ms and 3.2 ms respectively. These values, although higher than for Dpb5, are still much lower than for factors mainly associated with slowly diffusing mRNA molecules such as Gbp2. This indicates that, while a small fraction of Mex67 could be present in slowly diffusing complexes, the majority of Mex67 is not bound to large mRNPs.

### Mex67 is dynamically associated with the nuclear pore complex

Mex67 is concentrated at NPCs under steady state conditions (see Figure 1A), where its main function - namely mRNA export - is performed. Therefore, we aimed to further characterize the interaction of Mex67 with the NPC. To determine the amounts of Mex67 and other mRNA export factors at the NPC, we applied a recently developed quantitative image analysis tool termed NuRIM (Nuclear Rim Intensity Measurement) (Rajoo et al., 2018; Vallotton et al., 2019). In NuRIM, the nuclear envelope is segmented based on a red fluorescent marker targeted to the lumen of the endoplasmic reticulum (DsRed-HDEL), which then allows for the precise quantification of GFP signals at the nuclear envelope (Figure 2A). NuRIM was applied to Mex67 and Dbp5 as well as the nuclear pore components Nup84, Nup159 and Gle1 (Figure 2B). We found that Mex67 is present at the NPC in an amount comparable to Nup84, which is present in 16 copies per NPC (Rajoo et al., 2018). In contrast, Dbp5 has a 1.5 times higher copy number at the NPC than Nup84 (Figure 2C), suggesting that at steady state ~ 24 Dbp5 molecules reside at the NPC. Interestingly, this corresponds to the summed levels of the two Dbp5-binding partners at the NPC, Gle1 (8 copies) and Nup159 (16 copies).

**Figure 2:**
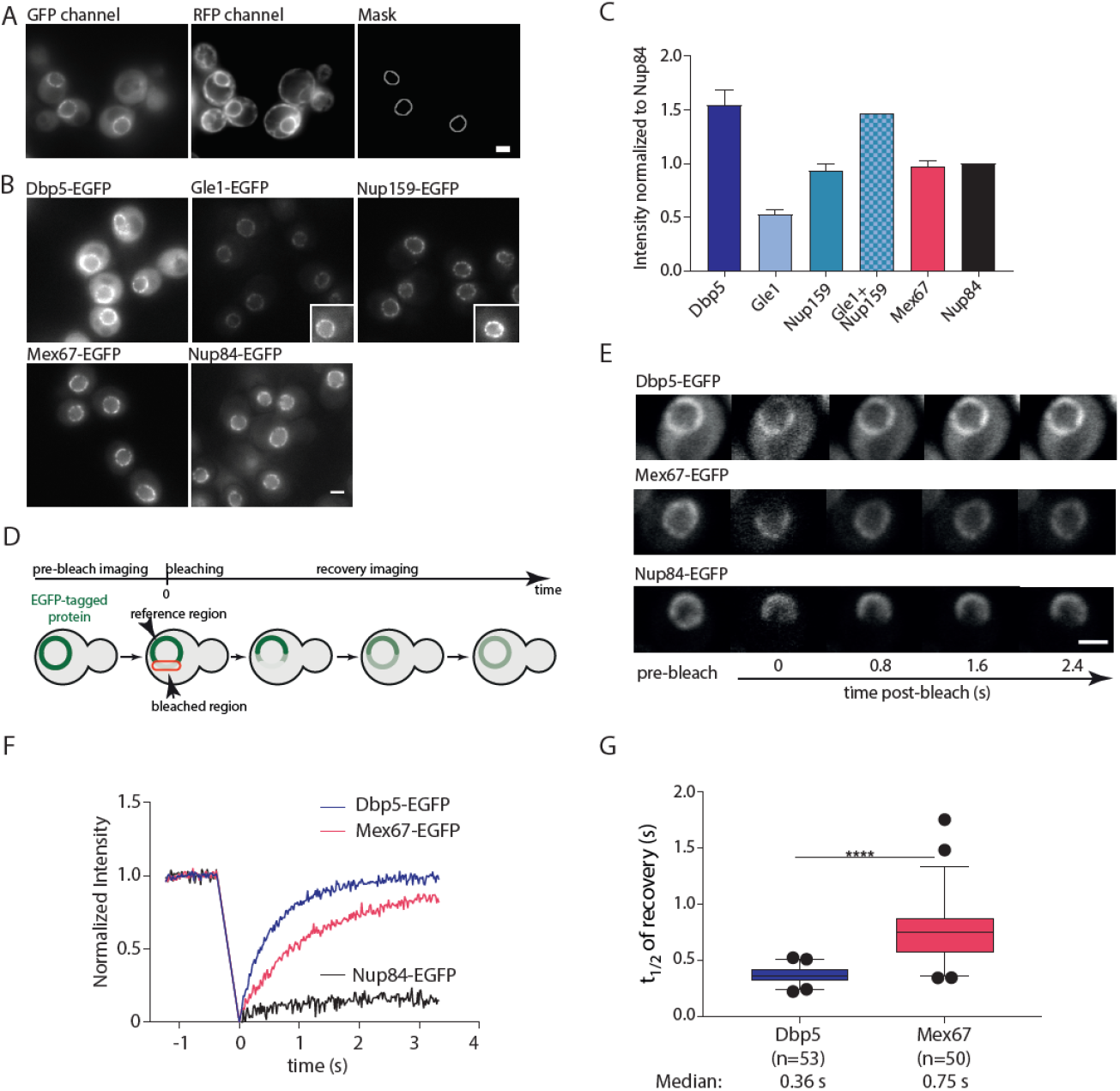
Mex67 and Dbp5 bind dynamically to the NPC. (A) Example for the production of the NuRIM analysis mask for quantification of GFP intensity on the nuclear envelope using DsRed-HDEL signal for nuclear envelope segmentation. (B) Representative fluorescence images of mRNA export factors and related Nups used for intensity analysis. Insets: example cells with increased contrast settings for better visualization. (C) Quantification of nuclear envelope intensities by NuRIM. Bars show the means of three biological replicates, the error bars represent the SEM. At least 1000 cells were analyzed per condition and replicate. (D) Schematic illustration of FRAP at the nuclear envelope. (E-G) FRAP of Dbp5, Mex67 and Nup84 tagged with GFP. (E) Representative FRAP images. (F) FRAP curves normalized to pre- and post-bleach values, mean curves of ≥ 50 cells are shown. (G) Half time of recovery retrieved from fitting individual FRAP curves with a single component model are plotted in boxplots (Line represents median, box represents 25-75 percentile, whiskers extend to 5-95 percentile. **** p value < 0.001 Mann-Whitney test). All scale bars 2 μm.

Next, we determined the dynamics of Mex67 and Dbp5 binding at the nuclear pore complex using fluorescent recovery after photobleaching (FRAP) (Figure 2D-G). Both Mex67 and Dbp5 quickly recovered within a few seconds after bleaching (Figure 2E-G). The median half time of recovery at the nuclear envelope was 0.36 s for Dbp5 and 0.75 s for Mex67 (Figure 2G). In contrast, the stable nucleoporin Nup84 showed no significant recovery in this timeframe (Figure 2F), indicating that NPC movement does not contribute to the measured dynamics. Thus, both Mex67 and Dbp5 are highly mobile at the NPC and are present in stoichiometries similar to nuclear pore proteins.

### Mex67 binding at the NPC does not depend on, but is modulated by mRNA and pre-ribosomes

Since Mex67 is thought to facilitate the passage of both mRNAs and pre-ribosomal subunits through the FG-repeat network of the NPC, we next asked whether the localization and dynamics of Mex67 is influenced by the presence of its cargo. To lower nuclear mRNA levels, we used the auxin-inducible degron (AID) (Nishimura et al., 2009) to acutely deplete the essential RNA Pol II subunit Rpb2. Auxin treatment led to a rapid and specific degradation of Rpb2, reduction in mRNA transcription and a concomitant loss of viability (Supplemental Figure 1A-B). Similarly, we inactivated RNA polymerase I to prevent the nuclear production of pre-ribosomes via auxin-induced degradation of the essential second largest subunit Rpa135. Depletion of Rpa135 led to reduced viability (Supplemental Figure 1C) as well as to reduced nucleolar size (Supplemental Figure 1D), indicative of impaired RNA Pol I function.

Neither inhibition of mRNA production by auxin-mediated depletion of Pol II, nor inhibition of ribosome production by depletion of Pol I led to a change in Mex67 abundance at the nuclear envelope (Figure 3A-D). This suggests that cargo is not required for Mex67 interaction with the NPC. However, we observed a small but significant increase in the turnover dynamics of Mex67 at the nuclear envelope in both mRNA and pre-ribosome depletion conditions (Figure 3E-J). Thus, while mRNA and ribosomal cargo do not influence the number of binding sites that are occupied by Mex67 at the NPC, the presence of cargo does appear to reduce the exchange rate of Mex67 at these binding sites. This suggests that cargo binding either increases the affinity of Mex67 to FG-repeats, e.g. by generating multivalency in the binding of Mex67, or reduces the mobility of Mex67 in the FG repeat network.

**Figure 3:**
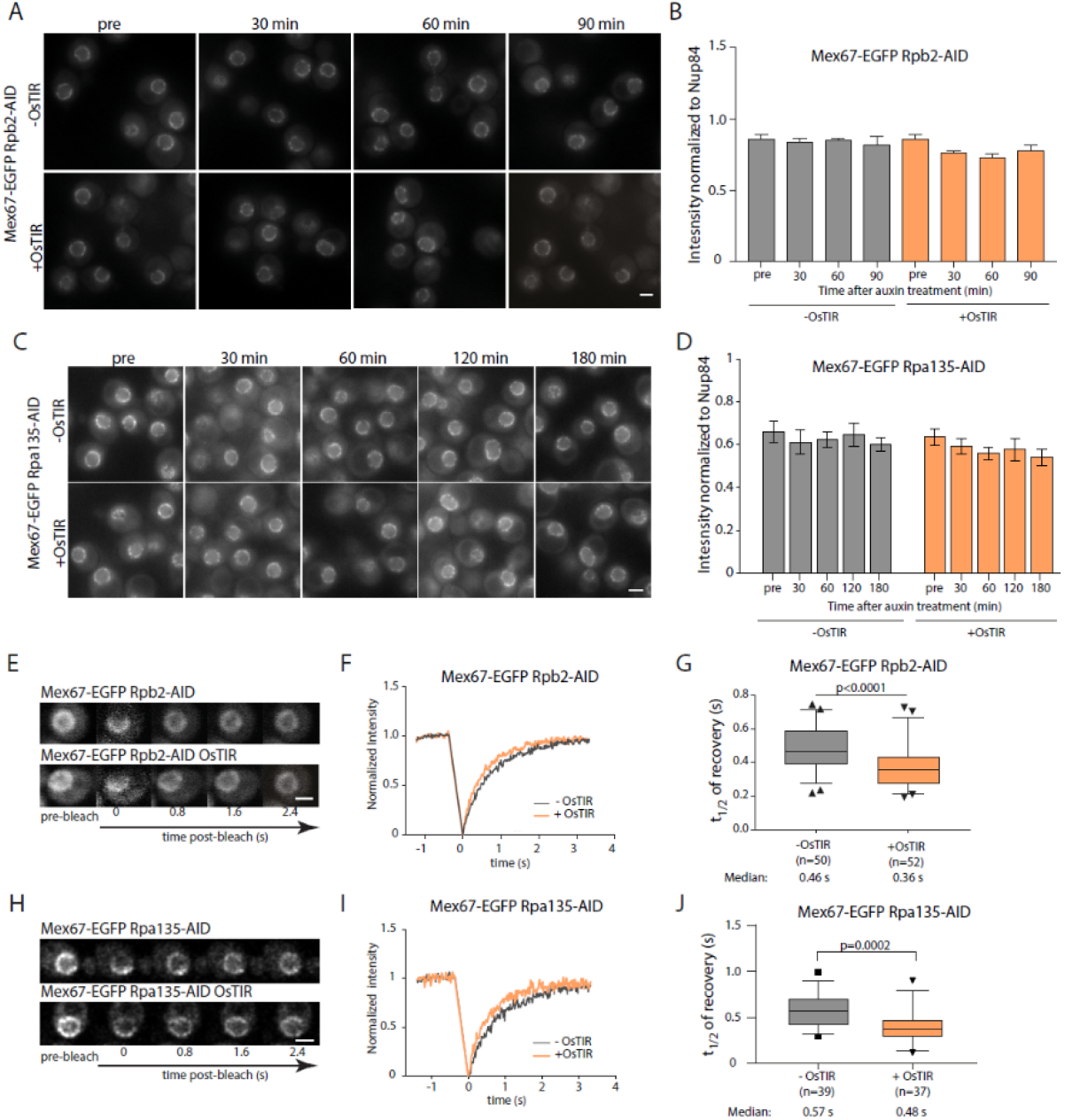
Mex67 dynamics, but not its amount at the NPC, is affected by the presence of cargo molecules. (A-D) NuRIM quantification of Mex67-EGFP intensity at the nuclear envelope upon depletion of RNA Pol II (A-B) or RNA Pol I (C-D) via auxin induced degradation. (A, C) Representative wide-field fluorescence microscopy images at different times after auxin treatment. (B, G) Intensity of Mex67-EGFP at the nuclear envelope. Mean of three biological replicates is shown, error bars represent SEM. At least 300 cells were analyzed per condition and replicate. (E-J) FRAP analysis of Mex67-EGFP dynamics at the nuclear envelope upon depletion of RNA Pol II or RNA Pol I via auxin induced degradation. (E, H) Representative images of FRAP experiments. (F, I) FRAP curves normalized to pre- and post-bleach values, mean curves of ≥ 50 cells are shown. (G, J) Half time of recovery retrieved from fitting FRAP. Line represents median, box represents 25-75 percentile, whiskers extend to 5-95 percentile, black dots represent data points outside this range. Data in F and G are pooled from three biological replicates, data in I and J from two. p values are from a Mann-Whitney test. All scale bars 2 μm.

### Mex67 binding at the NPC is not affected by Dbp5

Dbp5 was proposed to remodel mRNPs at the cytoplasmic side of the NPC and to remove Mex67 and other mRNA export factors from mRNA via its ATPase activity (Cole and Scarcelli, 2006a). To test whether ATP hydrolysis by Dbp5 affects Mex67 dynamics at the NPC, we depleted either Dbp5 itself or Gle1, which greatly stimulates ATP hydrolysis by Dbp5, using the AID system. Surprisingly, depletion of Dpb5 (Supplemental Figure 2A-B) did neither affect the steady-state levels of Mex67 at the NPC (Figure 4A-B) nor its turnover (Figure 4C-E). Depletion of Gle1, which led to a reduction of Dbp5 at the NPC (Supplemental Figure 2C) had also no effect on Mex67 intensity and dynamics at the NPC (Supplemental Figure 2D & E). These results argue that the interaction of Mex67 with the FG-repeat network of the NPC is not regulated by the activity of Dbp5. This is incompatible with the hypothesis that the removal of mRNA export factors from the RNA by Dbp5 releases these factors into the cytoplasm (Cole and Scarcelli, 2006a). Thus, while Dbp5 may decrease the interaction of Mex67 with mRNA, it does not influence its interaction with the NPC.

**Figure 4:**
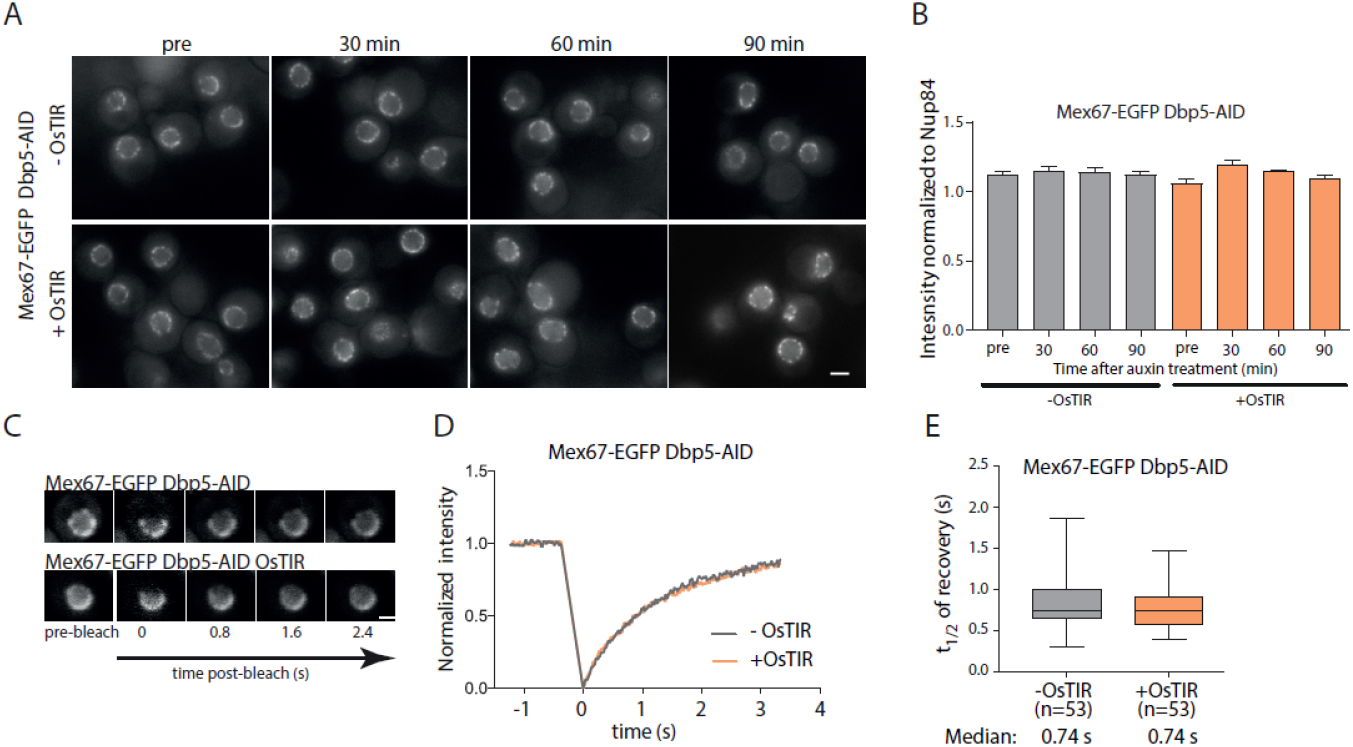
Mex67 binding to the NPC is independent of Dbp5. (A-B) NuRIM quantification of Mex67-EGFP intensity at the nuclear envelope upon depletion of Dbp5 via auxin induced degradation. (A) Representative wide-field fluorescence microscopy images at different times after auxin treatment. (B) Intensity of Mex67-EGFP at the nuclear envelope. Mean of three biological replicates is shown, error bars represent SEM. At least 300 cells were analyzed per condition and replicate. (C) FRAP analysis of Mex67-EGFP dynamics at the nuclear envelope upon depletion of Dbp5 via auxin induced degradation. (C) Representative images of FRAP experiments. (D) FRAP curves normalized to pre- and post-bleach values, mean curves of ≥ 50 cells are shown. (E) Half time of recovery retrieved from fitting individual FRAP curves. Line represents median, box represents 25-75 percentile, whiskers extend to 5-95 percentile. Data in D and E are pooled from three biological replicates. All scale bars 2 μm.

### The localization of Mex67 at the NPC is mediated by multiple FG repeats

As binding of Mex67 to the NPC does not depend on the presence of mRNA and is not modulated by Dbp5, we wanted to further characterize the FG-binding sites of Mex67 at the nuclear pore complex by applying NuRIM in a panel of FG repeat deletion mutants (Strawn et al., 2004, Supplemental Figure 3A). First, we examined GLFG deletions of Nups localized in the center of the NPC. The deletion of the adhesive GLFG repeats of either Nup49 or Nup57 alone or the combination of Nup57, Nup100 and Nup145 repeats had only moderate effects on Mex67 localization, reducing its level at the nuclear envelope by ~10 %. By contrast, deletion of the GLFG repeats of Nup116 lead to a more severe reduction, with a loss of ~ 20 % of Mex67 from the NPC. In addition, an increased pool of nucleoplasmic Mex67 could be detected in a subset of *nup116ΔGLFG* cells (Figure 5A&B). Next, we analyzed the effect of deleting FG repeat regions of asymmetrically localized Nups. While the deletion of the FG repeats of either Nup42 or Nup159 alone had no effect on Mex67 intensity at the nuclear envelope, the co-deletion of both these FG repeats on the cytoplasmic face of the NPC (Δcyt) led to a severe displacement of Mex67 from the NPC (reduction by approximately 40 %) and increased levels of Mex67 in the nuclear interior. Both nucleoplasmic enrichment and loss from the nuclear envelope were heterogenous phenotypes and varied in different cells (Figure 5A). A similar severe mislocalization of Mex67 was also observed in a triple deletion of the FxFG repeats of the three nuclear basket Nups, Nup1, Nup2 and Nup60 (Δnuc) which was exacerbated in cells where the deletion of cytoplasmic and nucleoplasmic FG repeats were combined (Δcyt+Δnuc). Note that nuclei with very low EGFP signal are excluded by the NuRIM algorithm, leading to an overestimation of the average intensity at the nuclear envelope in these severely compromised strains (Figure 5B). Overall, our results suggest that Mex67 does not have a single, predominant NPC interaction site but instead multiple FG-repeats contribute, directly or indirectly, to the steady-state localization of Mex67 at the NPC.

**Figure 5:**
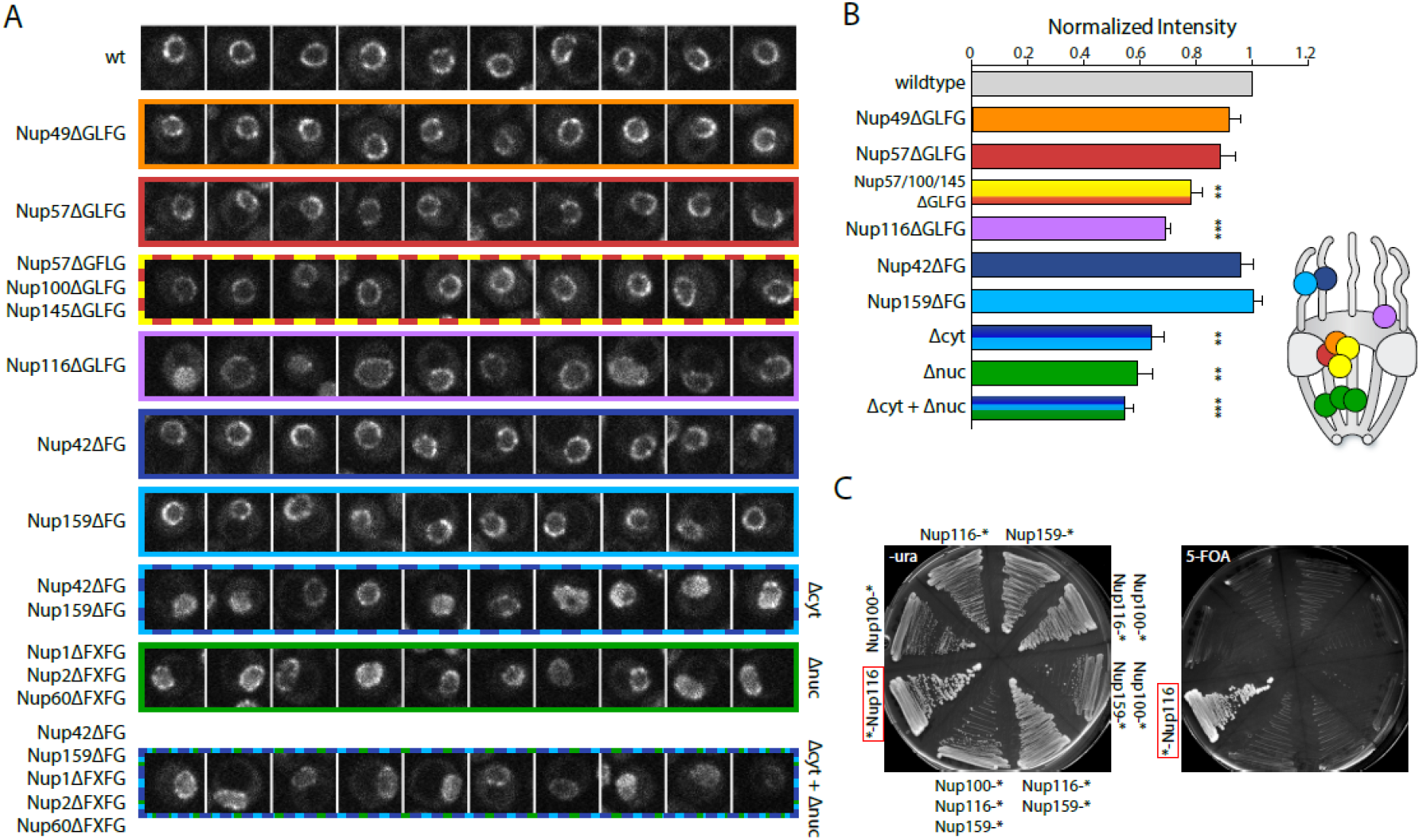
Mex67 relies on multiple FG repeats for binding the NPC and fulfills its essential function only at the NPC. (A) Example nuclei are shown for different strains expressing Nups that have deleted FG/GFLG/FXFG regions as indicated. Images are single slices of z-stacks acquired on a spinning disk microscope. Slices were selected for being equatorial planes using an ER marker channel. Scale bar 2 μm. (B) Intensity of Mex67 at the nuclear envelope using NuRIM in different strains with deletions of various FG repeats. Shown is the mean of three biological replicates normalized to wildtype cells. Error bars represent the SEM. Stars indicate significant p values of a paired t-test compared to wild type (** p<0.01, *** p<0.001). Schematic indicates approximate positions of the NPC components in the NPC structure. Note that this applies to the body of the protein and not necessarily to its FG repeat region. (C) N-terminal fusion of Mex67 to Nup116 rescues deletion of *MEX67* locus. Strains in which a deletion of the *MEX67* locus was covered with Mex67 expressed from a plasmid containing the *URA3* selection marker gene were engineered to express fusions of Mex67 to different nucleoporins. Streaks of these strains on 5-FOA containing plate indicate rescue only by the N-terminal Nup116 fusion. * indicates the position of Mex67 fusion.

### The essential function of Mex67 is restricted to the NPC

Our results so far do not support a model where Mex67 functions as a shuttling transport receptor that binds to its substrate in the nucleus and is released into the cytoplasm at the termination of export. Instead, they suggest that Mex67 behaves like a dynamic nuclear pore component, where it resides in the FG-repeat network independently of mRNA. We therefore tested whether the essential function of Mex67 could be entirely restricted to the nuclear pore complex, and asked whether a protein fusion of Mex67 to a nucleoporin could rescue a deletion of the endogenous *MEX67* gene. To this end, we generated strains where Mex67 was fused to different components of the nuclear pore complex in a background where a deletion of *MEX67* was covered by a plasmid carrying the full *MEX67* locus as well as the *URA3* selection marker (Segref et al., 1997). Since the deletion of the GLFG repeats of Nup116 as a single component had the most drastic effect on Mex67 localization, we generated N- as well as C-terminal fusions of Mex67 to Nup116. In addition, C-terminal fusions with Nup100, Nup1 and Nup159 were tested. Nup-Mex67 fusion strains were tested for growth on 5-FOA (5-fluoroorotic acid) plates to identify strains that are viable in the absence of the wild-type Mex67 protein (Figure 5C). Remarkably, the N-terminal fusion of Mex67 to Nup116 supported growth on 5-FOA, suggesting that Mex67 is able to perform its essential function even when it is permanently tethered to the NPC via Nup116. After loss of the Mex67 plasmid, the Mex67-Nup116 strain exhibited only slight growth defects when compared to the parental strain at 30 °C or 37 °C although it showed delayed growth at lower temperatures (Supplemental Figure 3B). Western blotting confirmed that the only Mex67 variant detectable in this strain is a large fusion protein (Supplemental Figure 3C). In addition, a C-terminal EGFP-tag on Mex67-Nup116 was used to verify the identity of the high molecular weight band as the Mex67-Nup116 fusion protein and to confirm the exclusive and stable localization of the fusion protein to the nuclear envelope (Supplemental Figure 3D-E). Fusion of Mex67 to the flexible FG repeat region of Nup116 thus allows it to perform its function even when stably tethered to the NPC, showing that its essential function is spatially restricted to the NPC.

### Conclusion

In summary, we have shown that the RNA export factor Mex67 functions as a mobile nuclear pore component that dynamically binds to the NPC independently of cargo. Furthermore, the interaction of Mex67 with the NPC is not modulated by the DEAD-box ATPase Dbp5. These findings are consistent with a recent report in mammalian cells (Ben-Yishay et al., 2019). Since Dbp5 is thought to impart directionality to the mRNA export process by releasing export factors form cargo, this suggests that the action of Dbp5 leads to the release of mRNA from Mex67 but not of Mex67 from the NPC. By fusing Mex67 to a nucleoporin, we further demonstrate that the essential function of Mex67 occurs entirely at the NPC and propose that Mex67 can receive its export cargo within the central channel of the NPC to facilitate its translocation across the FG-repeat diffusion barrier. In this model, loading of the mature mRNP onto Mex67-Mtr2 happens at the nuclear pore complex. It will now be interesting to further determine, which steps of mRNP maturation occur in the nucleoplasm and which processes are performed at the nuclear pore complex.

## Materials and Methods

### Plasmid construction

Plasmids were constructed using standard molecular biology techniques. Fragments generated by PCR were verified by sequencing. pKW3124 was clones by restriction cloning inserting a Nop1-RFP fragment from pRS416-NOP1-RFP (Rothstein lab) into pRS406 with NotI/SalI. In pKW3235 and pKW3570, the A206->K206 mutation was introduced to obtain monomeric GFP (Zacharias et al., 2002). pKW4216 used for mating type switching in URA+ strains was derived from YCp50-HO (Russell et al., 1986). pKW4547 was constructed by insertion of Mex67 into a pFA6a-hisMX6 plasmid with a long N-terminal linker of 110 amino acids. pKW4648 was constructed by replacing EGFP in pKW4588 (Vallotton et al., 2019) with Mex67 using RecA cloning and changing the selection marker from URA3 to LEU2 using restriction cloning. pKW4688 was derived from pKW4648 by addition of a segment of the Nup116 promoter upstream of the selection marker to allow for single integration. All plasmids used in this study are listed in Supplemental Table 1. Primers are listed in Supplemental Table 2.

### Yeast strain construction

*S. cerevisiae* strains were constructed using standard yeast genetic techniques either by transformation of a linearized plasmid or of a PCR amplification product with homology to the target site (Baudin et al., 1993). Strains used for FCS were tagged with a monomeric GFP version obtained by the A206->K206 mutation. Strains with a deletion of *MEX67* covered by shuffle plasmid were based on the Mex67 shuffle strain received from the Hurt group (Segref et al., 1997). Strains with different deletions of FG repeats were based on strains received from the Wente group (Strawn et al., 2004) by integration of an ER marker (DsRed-HDEL) and tagging of Mex67 with EGFP. All yeast strains are listed in Supplemental Table 3.

### Yeast culture conditions

Cells were cultured in YPD (for Western Blots, growth assays) or complete synthetic medium supplemented with extra adenine (for microscopy) in the presence of 2 % glucose. Cells were grown at 25 °C for most experiments. All experiments were carried out with cells in exponential growth phase. For microscopy experiments, cells were usually inoculated from saturated cultures into fresh medium and grown overnight to OD_600_ 0.3-0.8 and then imaged. Alternatively, cells were diluted in the morning and grown for additional 2-3 cell cycles before the start of the experiment. Auxin treatment was performed with auxin (Indole-3-acetic acid, 500 μM, stock 500 mM in ethanol) and IP6 (phytic acid dipotassium salt, 4 μM, stock 40 mM in water). As a solvent control, cells were treated with IP6 and ethanol.

### Microscopy

Cells were pre-grown and inoculated in 1 ml in 24 well plates overnight so that they reached an OD of 0.3-0.8 for the experiment. Cells were then transferred to 384 well plates (Matrical) or 8-well Ibidi dishes coated with concanavalin A (stock solution of 0.2 mg/ml, air dried).

Images for Figure 5A were acquired on a SpinningDisk microscope (Yokogawa Confocal Scanner Unit CSU-W1-T2) built on a Nikon TiE body and controlled with the VisiVIEW software and a 100x NA 1.49 CFI Apo TIRF objective. The camera was an iXon Ultra EMCCD (Andor), excitation lasers were a DPSS 488 nm (200 mW) and a diode 561 nm (200 mW) laser. Z scanning was performed in streaming mode with a LUDL BioPrecision2 Piezo Stage with 100 ms exposure times per frame. Filters were: Dichroic quad-band DAPI/GFP/RFP/CY5, splitting filter to camera ports: 561LP, emission filters GFP/ET525/50 and mCherry ET630/75 respectively. Imaging for Figure 6A was performed on the same system in widefield-mode with excitation from Spectra X LED lines at 475 nm and 542 nm.

### Western Blotting

Cells were grown overnight at 25 °C and diluted to an OD_600_ of 0.2. Cells were grown for additional 2-3 cell cycles at 25 °C shaking until they reached an OD_600_ of 0.6-0.9. A volume of yeast culture of was spun down, supernatant was discarded and cells were incubated in 1 ml of 1 M NaOH for 10 min. Afterwards cells were spun down and resuspended in 30-60 μl 1× SDS buffer/DTT 100 mM, boiled for 5 min at 95 °C and stored at −20 °C. The samples were thawed, boiled again for 5 min at 95 °C. The samples were usually loaded on 12% resolving SDS PAGE gel or, in case of the fusions Mex67-Nup116, on a precast NuPAGE™ 4-12 % Bis-Tris Protein Gels (Invitrogen). Sample loading was adjusted based on equivalent OD_600_ in each lane. The SDS PAGE was run for 50 - 60 min at 200 V. The proteins were blotted onto a nitrocellulose membrane (Amersham Protran 0.2 NC, GE Healthcare life sciences) for 1-2h at 300 mA. Blots were blocked in 5% milk in TBST for at least 2h or overnight. The first antibody was applied overnight in 1-5% milk in TBST on rotating wheel at 4 °C. The membrane was washed three times in TBST for 20 min and stained with the secondary antibody in 1-5% milk in TBST for 40 min at RT. The blot was washed with TBST three times for 20-40 min before imaging with the CLx ODYSSEY Li-COR. Antibodies used were: rabbit anti-Mex67 (Dargemont Lab, Diderot University, Paris, 1:2000), mouse monoclonal anti-GFP (Roche, 11814460 001, 1:2000), monoclonal mouse anti V5 (Invitrogen R960-25, 1:2000), mouse monoclonal anti-yeast PGK (22C5D8, Thermo Fisher Scientific, 4592250, 1:3000), goat anti mouse IgG-Alexa 680 (Thermo Fisher Scientific, A-21057, 1:10000) and goat anti-rabbit IgG IRDye800CW (Li-COR Biosciences, 926-32211, 1:10000).

### NuRIM fluorescence intensity measurements

NuRIM fluorescence intensity quantification experiments were carried out on a temperature controlled Nikon Ti Eclipse equipped with a Spectra X LED lamp using a Apochromat VC 100× objective NA 1.4 (Nikon) (filters: Spectra emission filters 475/28 & 542/27 and DAPI/FITC/Cy3/Cy5 Quad HC Filterset with 410/504/582/669 HC Quad dichroic and a 440/521/607/700 HC quadband filter (Semrock)) with exposure times of 500 ms in the GFP and 1 s in the RFP channel. 30 image frames were imaged per sample and condition.

Images were processed using NuRIM code (Rajoo et al., 2018; Vallotton et al., 2019) in MATLAB® (MathWorks®). Briefly, this method uses the HDEL-DsRed signal to generate a mask that segments the nuclear envelope and then extracts intensity from this mask. Machine learning is applied to remove the contribution from variable background signals.

### Fluorescence recovery after photobleaching

FRAP experiments were performed on the Leica TCS SP8 STED/FLIM/FCS or a Leica TCS SP8-AOBS microscope using a 63× 1.4NA Oil HC PL APO CS2 objective. Bidirectional scanner at speed of 8000Hz, NF488/561/633, an AU of 1.5 and a FRAP booster for bleaching were applied for every FRAP experiment using the PMT3 (500-551nm) and PMT5 (575-694nm) detectors. Image size of 504 × 50 and a zoom of 3.5 were used together with line accumulation of three, yielding a frame rate of 60.61 frames / s with a pixel dwell time of 73 ns. 50 pre-bleach and 200 post-bleach frames were acquired. A 488 nm argon laser line was used at 50 % power in addition to a 561 nm DPSS laser line. The bleach was performed using 100% argon laser power for 200 ms. Imaging was conducted with 1.5% laser intensity with a gain of 800 to illuminate the GFP, and 0.3% of the 561 laser power to illuminate the HDEL with DsRed, was used during pre- and post-bleach acquisition. The photobleaching was performed by manually defining an elliptical region comprising approximately one-third of the cell nucleus. The mobility of GFP-labeled proteins in the bleached NE region was evaluated by quantifying the signal recovery in the bleached region. Extracellular background (Ibg) was subtracted from the intensity of the bleached region (I_bl_) and the values were bleach-corrected by normalizing for total cell intensity (I_total_) resulting in (I_bl_-I_bg_)/(I_total_-I_bg_) (Bancaud et al., 2010). The datasets acquired during three independent experiments were processed using MATLAB with custom written scripts. Briefly, the intensities were normalized for pre and post bleach values and the curves fitted with an exponential function ft= A.*(1−exp(−r.*t) (ft= normalized fluorescence intensity, A = non-recovering pool, r= time constant of recovery, t = time) with the method “NonlinearLeastSquares”. The measurements of three independent experimental days were pooled. Plots were generated in GraphPad Prism 7 (GraphPad).

### Fluorescence correlation spectroscopy FCS

FCS experiments were performed with a Leica TCS SP8 STED/FLIM/FCS microscope equipped with a 63× 1.2NA W HC PL APO CS2 objective with the HyD 2 SMD (Single Molecule Detection) (500-550nm) and HyD 4 SMD (604-750nm). The Leica components were used in combination with the PicoHarp 300, TCSPC Module (Time Correlated Single Photon Counting system), 4 Channel Detector Router (PHR 800) and the Software SymPho Time 64 1.6. FCS experiments were conducted using the 488 nm Argon laser line at 20% power set to 0.1-0.2 % for acquisition and the pulsed white light laser (WLL) at 70% with a laser pulse of 20 MHz at 594 nm set to 0.3-0.7 % for acquisition. Cells were imaged for 30 s and traces were recorded and saved by the PicoQuant software. FCS measurements were performed in diploid yeast cells in the center of the nucleus in order to place the entire confocal volume with high confidence within the nuclear boundary.

Analysis of FCS data was performed using Fluctuation 4G analyzer (Wachsmuth et al., 2015). Raw photon traces with strong fluctuations were discarded. Auto-correlation curves of the accepted in vivo photon traces were calculated over a rolling time window of 0.5 to 5 s and averaged over the whole duration of the photon trace (Wachsmuth et al., 2015). This approach allows removing the contribution of slow fluctuations arising from photobleaching or small cell movement. Auto-correlation curves were fit with a one-component anomalous di**ffusion model includi** ng the fluorescence blinking contribution. This model gave access to a mean diffusion time corresponding to the average duration of the transit of the fluorescent molecule through the confocal volume. When fitting the auto-correlation curves, the lifetime of the dark state was fixed to 100 μs (Wachsmuth et al., 2015) and the structural parameter kappa was fixed to 5, based on experimentally determined values. Anomality parameters obtained from fits for EGFP, Dbp5 and Mex67 were 0.84 ± 0.04, 0.80 ± 0.02 and 0.80 ± 0.01 in the nucleus and 0.98 ± 0.03, 0.71 ± 0.01 and 0.84 ± 0.01 in the cytoplasm respectively (mean ± standard error of the mean (SEM)).

### Fluorescence *in situ* hybridization

Fluorescence *in situ* hybridization was carried out according to (Dultz et al., 2018). Briefly, cells were grown in to exponential growth phase at 30 °C. Cells were then treated with auxin and IP6 and fixed after one (Dpb5-AID) or two (Rpb2-AID) hours for 25 min at 30 °C and with 4 % paraformaldehyde (EM grade 32 % paraformaldehyde acqueous solution electron Microscopy Sciences 15714), washed with buffer B (1.2 M sorbitol, 100 mM KHPO4 at pH 7.5, 4 °C) and stored at 4 °C overnight. Cells were then spheroplasted for 20 min using 1 % 20T zymolyase in 1.2 M sorbitol, 100 mM KHPO4 at pH 7.5, 20 mM vanadyl ribonuclease complex and 20 μM β-mercaptoethanol, washed with buffer B to stop the spheroplasting reaction and then washed into 10 % formamide (Merck Millipore S4117) in 2× SSC.

DNA probes targeting *FBA1* coupled to CalFluore Red (Stellaris, LGC Biosearch, Novato, CA; probes were synthesized by BioCat, Heidelberg, Germany, Supplemental Table 4) or oligo dT30 coupled to ATTO488 (Microsynth) were used. Per sample, 0.5 μl of *FBA1* probe mix (stock 25 μM) or 0.25 μl of oligo dT30 (stock 50 μM) was mixed with 2 μl of salmon-sperm DNA (10 mg/ml, Life Technologies, 15632-011) and 2 μl yeast transfer RNA (10 mg/ml, Life Technologies, AM7119). The probe mix was denatured in 50 μl per sample of hybridization buffer F (20 % formamide, 10 mM NaHPO_4_ at pH 7.0) for 3 min at 95 °C and then mixed with 50 μl per sample hybridization buffer H (4× SSC, 4 mg/ml BSA (acetylated) and 20 mM vanadyl ribonuclease complex). Cells (approximately corresponding to the pellet of 5 ml initial culture) were resuspended in the hybridization mix and incubated overnight at 37 °C. After four washing steps (10 % formamide/2× SSC; 0.1 % Triton/2× SSC; 2× SSC/DAPI; 2× SSC), cells were stored at 4 °C. Cells were imaged in concanavaline A coated 384 wells. Microscopy was performed with an inverted epi-fluorescence microscope (Nikon Ti) equipped with a Spectra X LED light source and a Hamamatsu Flash 4.0 sCMOS camera using a PlanApo 100 × NA 1.4 oil-immersion objective and the NIS Elements software. 27 z slices were acquired at 200 nm spacing.

## Acknowledgements

We thank Malthe Wachsmuth for his help with Fluctuation Analyzer 4G for FCS data analyses. We are grateful to Justine Kusch from ScopeM, ETH Zürich, for microscopy support, and to Catherine Dargemont for providing the anti-Mex67 antibody. We thank members of the Weis group for discussions and critical reading of the manuscript. We are particularly grateful to Stephanie Heinrich for helpful discussions and advice. Support from the Swiss National Science Foundation (project number: 31003A_179275) is acknowledged by KW. The authors declare no competing financial interests.

## Author Contributions

This work was devised and planned by CPD, ED and KW. Experiments were carried out by CPD, ED, RM and PV. Data were analyzed by CPD, ED and PV. Code for data analysis was produced by PV and ED. The FCS experiments were supported by SH. The manuscript was written by CPD, ED and KW and revised by all authors.

**Abbreviations**
AID: auxin-inducible degron
NPC: nuclear pore complex
SEM: standard error of the mean
FRAP: fluorescence recovery after photobleaching
FCS: fluorescence correlation spectroscopy

---

**Supplemental Figure 1:**
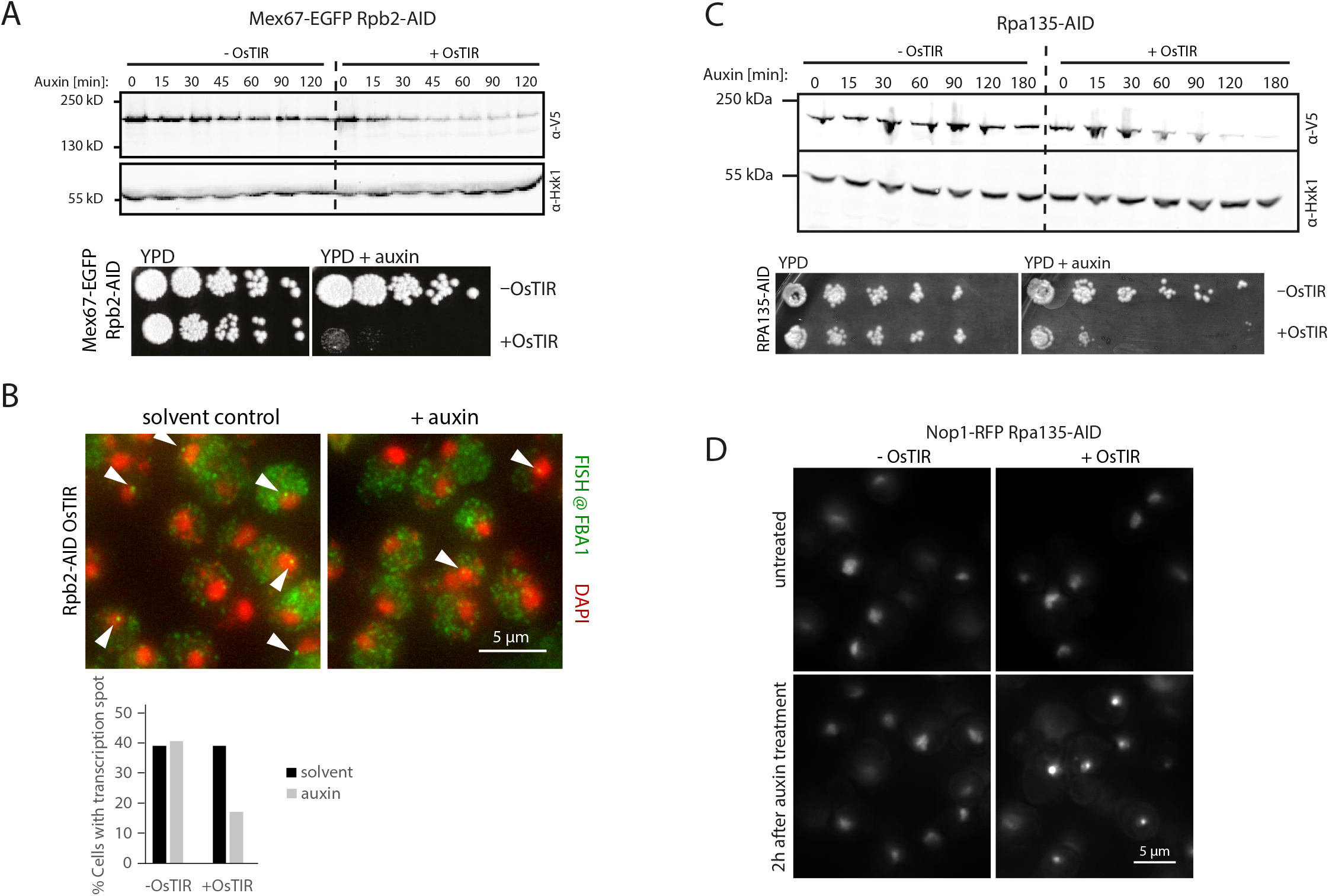
AID-mediated depletion of RNA Pol II and RNA Pol I. (A) Rpb2 is degraded after auxin treatment. Top: Lysate from cells treated with auxin were analyzed for presence of Rpb2-V5-AID. Bottom: Depletion of Rpb2 rendered cells inviable. (B) Depletion of Rpb2 leads to loss of transcription. FBA1 mRNA was visualized by single molecule fluorescence in situ hybridization (smFISH). Transcription foci, i.e. bright dots of multiple nascent mRNAs, are observed in the nucleus (arrowhead) of 40 % of control cells but only in ~ 17% of cells treated with auxin for 1 hour (one of two biological replicates shown). (C) Rpa135 is degraded after auxin treatment. Top: Lysate from cells treated with auxin were analyzed for presence of Rpa135-V5-AID. Bottom: Depletion of Rpa135 rendered cells inviable. (D) Depletion of Rpa135-AID leads to reduction of nucleolar size as visualized by the nucleolar marker Nop1-RFP.

**Supplemental Figure 2:**
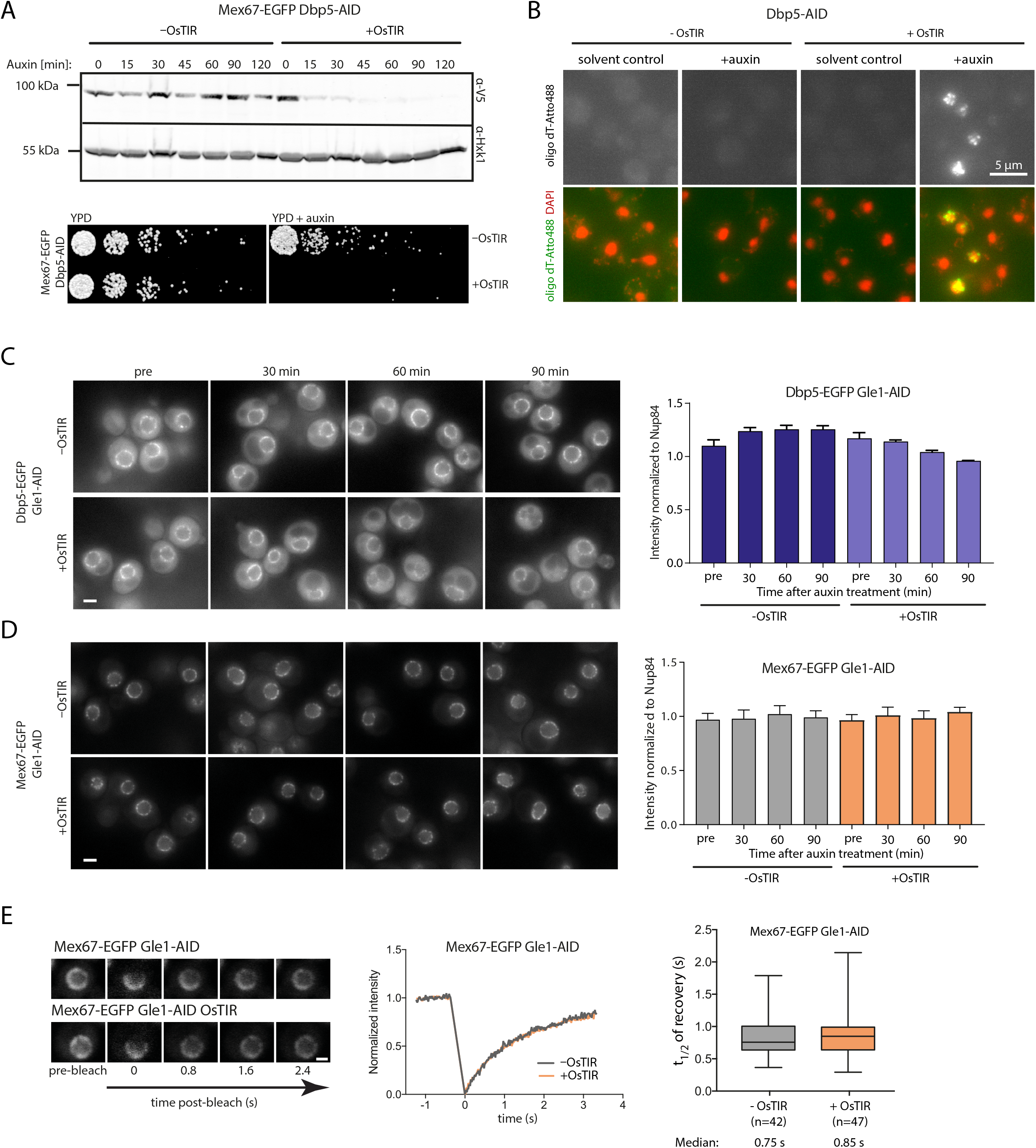
Depletion of Dbp5 and Gle1 via an auxin inducible degron. (A) Dbp5 is effi ciently degraded after auxin treatment. Top: Lysate from cells treated with auxin were analyzed for presence of Dbp5-V5-AID. In cells expressing OsTIR, Dbp5 was effi ciently depleted within 30 min. Bottom: Depletion of Dbp5 rendered cells inviable. (B) Oligo dT signal accumulates in nuclei of cells depleted for Dbp5 using an auxin inducible degron. Yeast cells expressing or not expressing OsTIR where treated with auxin or ethanol as solvent control for one hour. Many cells depleted for Dbp5 showed accumulation of oligo dT signal in puncta in the nucleus. Images shown represent maximum projections. (C-D) Representative images and NuRIM quantification of Dbp5-EGFP (B) and Mex67-EGFP upon auxin-induced depletion of Gle1. Scale bar 2 μm. Bars show the means of three biological replicates, the error bars represent the SEM. In each experiment, at least 300 cells were analyzed per condition. (E) FRAP of Mex67-EGFP upon depletion of Gle1. Left: Representative FRAP images. Scale bar 2 μm. Middle: FRAP curves normalized to pre- and post-bleach values, mean curves of ≥ 50 cells are shown. Right: half time of recovery retrieved from fitting individual FRAP curves. Line represents median, box represents 25-75 percentile, whiskers extend to 5-95 percentile.

**Supplemental Figure 3:**
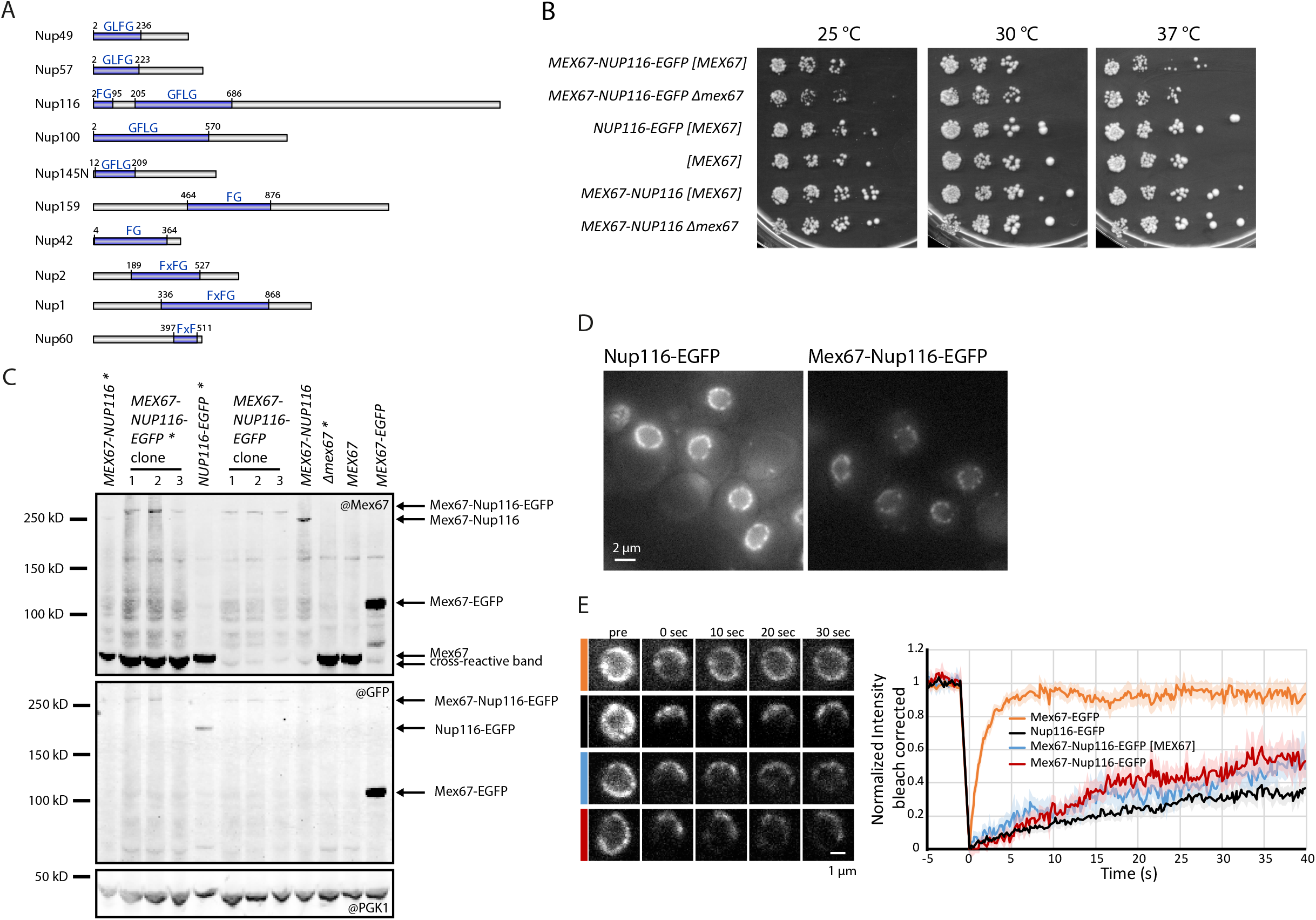
Mex67-Nup116 can replace wildtype Mex67. (A) Schematic representation of truncated nucleoporins expressed in strains used in Figure 5. Deleted repeat regions are shown in blue with the position of the first and last deleted residue inidcated. Blue labels name the type of repeats predominantly found in the deleted domains. In some nucleoporins, a small number of repeats can still be present in the intact regions of the proteins. For a complete description and depiction of the deletion mutants compare Strawn et al. 2004. (B) Spot assays shows minor growth defect and slight cold sensitivity of Mex67-Nup116 fusion strains after loss of [MEX67] plasmid. Cells were plated at 1:5 dilutions onto YPD plates and grown at the indicated temperature for 3 days. (C) Western blot showing expression Mex67-Nup116 fusion constructs. * indicates presence of [MEX67] plasmid. (D) Wide-field fluorescence images of Nup116-EGFP and Mex67-Nup116-EGFP. Contrast adjusted identical for the two images. (E) FRAP analysis of Mex67-Nup116-EGFP dynamics at the nuclear envelope. Left: Representative images of FRAP experiments Right: FRAP curves normalized to pre- and post-bleach values, mean curves of 20 cells are shown.

